# Multiomic profiling reveals aberrant immunomodulatory signature in β-propeller protein-associated neurodegeneration patient iPSC-derived microglia

**DOI:** 10.1101/2025.09.04.668126

**Authors:** Gamze Ӧzata, Rachel M. Wise, Aida Cardona-Alberich, Naiyareen F. Mayeen, Stephan A. Müller, Stefan F. Lichtenthaler, Luigi Zecca, Lena F. Burbulla

**Author notes:** Corresponding author: Lena Burbulla, PhD, Professor for Metabolic Biochemistry of Neurodegenerative Diseases, Biomedical Center (BMC), Faculty of Medicine, Ludwig-Maximilians-Universität, München & German Center for Neurodegenerative Diseases (DZNE) Munich. Contributed equally (Gamze Ӧzata, Rachel M. Wise).

## Abstract

Microglia are the primary immune cells of the central nervous system and play a crucial role in maintaining brain homeostasis. In common neurodegenerative diseases such as Alzheimer’s disease and Parkinson’s disease (PD), early and sustained microglial activation has been shown to precede neuronal loss, with elevated levels of microglia-derived inflammatory mediators detected in affected brain regions. In contrast, little is known about the role of microglia in rare neurodegenerative disorders. One such disorder is β-propeller protein-associated neurodegeneration (BPAN), a common subtype of neurodegeneration with brain iron accumulation (NBIA). BPAN shares pathological features with PD, including iron accumulation and selective loss of dopaminergic neurons in the substantia nigra, and is caused by mutations in the *WD repeat domain 45 (WDR45)* gene encoding an autophagy protein also called WIPI4. However, the pathological role of mutant WDR45 in BPAN and the possible contribution of microglia remain unresolved.

We generated the first BPAN patient microglia model system using induced pluripotent stem cells (iPSCs) to identify immune-related alterations and immunomodulatory signaling changes in a disease-relevant context. Integrated transcriptomic and proteomic profiling of iPSC-derived microglia from BPAN patients revealed a consistent shift from a homeostatic to a reactive, disease-associated state. Transcriptomic analysis showed disruption of core microglial pathways, including immune activation, stress response, and autophagy, consistent with a chronic pro-inflammatory phenotype. Complementary secretome analysis identified impaired lysosomal function and increased antigen presentation pathways, further supporting persistent microglial activation. Together this suggests that dysfunctional microglial states may contribute to BPAN pathogenesis. Our findings lay the groundwork for advancing immunomodulatory research in BPAN and may open new avenues for therapeutic development targeting microglial dysfunction.

## Introduction

β-Propeller protein-associated neurodegeneration (BPAN) is a subtype of neurodegeneration with brain iron accumulation (NBIA), a family of ultra-rare neurological disorders^1–3^. This disorder is caused by primarily *de novo* mutation of the X-linked *WD repeat domain 45* (*WDR45*) gene which encodes the protein WDR45 (also called WIPI4) implicated in autophagy^4–6^. Analogous to autism spectrum and Rett syndrome-like disorders, pediatric BPAN patients often display epilepsy, developmental delays, cognitive impairment, and behavioral and language deficits^7^. The adolescent/adult phase is characterized by the onset of parkinsonism, dystonia, and dementia, evocative of Parkinson’s disease (PD) and Alzheimer’s disease (AD). Moreover, cellular-level pathological phenotypes – including diffuse tau-positive neurofibrillary tangles^8^, thinning of the corpus callosum^9,10^, abnormal excess iron accumulation, and severe atrophy of dopaminergic neurons in the substantia nigra pars compacta (SNpc), suggest that BPAN may also share molecular mechanisms with these more common neurodegenerative diseases^2,8^.

Microglia are the resident immune cells of the central nervous system (CNS) critical for maintaining brain health and homeostasis through the complex orchestration of immune surveillance, synaptic pruning, debris clearance, and rapid responses to changes in the neuronal environment. In neurodegenerative diseases, microglia undergo dynamic transformations from surveillant to activated or “reactive” states, marked by the production of pro-inflammatory cytokines, altered phagocytic function, and increased antigen presentation. This shift can be protective in early stages, but when sustained, may promote chronic neuroinflammation and neurodegeneration. Indeed, accumulating evidence from several neurodegenerative conditions has implicated dysregulated microglial function as an early and central player in disease progression, rather than a secondary response to neuronal death^11,12^. However, the role of microglia in rare neurodegenerative diseases remains underexplored.

Intriguingly, the SNpc contains one of the highest densities of microglia in the brain^13^, which react to aggregated forms of the PD-associated protein α-synuclein^14^ as well as neuromelanin-laden apoptotic neurons^15^ by releasing cytokines, chemokines, and other signaling molecules including reactive oxygen species (ROS), nitric oxide (NO), and complement factors^11^. In PD patient brains, excessive microglial activation precedes dopaminergic neuron death, and elevated microglial-derived inflammatory mediators have been observed in the SNpc, suggesting that reactive microglia may contribute to the preferential demise of these vulnerable neurons^16–24^. Most importantly, it has been shown that autophagy regulates microglial activation by inhibiting pro-inflammatory activity in resting conditions^25,26^, which makes the process of autophagy a key mediator of microglial-driven inflammation. Reports have shown that impaired microglial autophagy correlates with increased neuroinflammation, neuronal iron dyshomeostasis, and dopaminergic neuron death^25,27–30^. *In vivo*, knockout of autophagy genes specifically in microglia results in elevated neuroinflammation and worsened degeneration of nigral dopaminergic neurons, even manifesting the PD-like symptoms of impaired motor function and cognitive decline^31,32^. Interestingly, evidence of neuroinflammation in BPAN patients^9,10,33^, and elevated gliosis in one *Wdr45-* knockout mouse model^34^ indicate that aberrant microglial activity may be another shared pathology with PD, yet their potential involvement in BPAN pathology remains unexplored. This is particularly important as animal models of dopaminergic degeneration have shown that depleting microglia^35^ or inhibiting their pro-inflammatory activation^36–38^ results in attenuated destruction of this vulnerable neuron population, thus microglial modulation may represent a novel target for early therapeutic intervention and protection of dopaminergic neurons.

*In vitro* systems^39–42^ have provided valuable insight into the role of WDR45 in autophagosome initiation and elongation, and transgenic mouse models^34,43,44^ have elucidated the pathological consequences of defective *Wdr45* on both autophagic machinery and function in mouse neurons. However, existing cellular models do not recapitulate the unique complexity of dopaminergic neuron physiology, and available animal models do not embody important species-specific differences between mouse and human nigral dopaminergic neurons^45^. Thus, without proper manifestation of disease-relevant phenotypes in human brain cells, the precise roles of mutant *WDR45* in BPAN pathogenesis remain elusive. Advancements in induced pluripotent stem cell (iPSC) technology have facilitated the investigation of human neurons, and more recently glial cells^46^, providing an inimitable platform to define pathogenic phenotypes, probe for underlying mechanisms, and screen rescue strategies in patient cells. Indeed, from the only two available reports using iPSC-derived dopaminergic neurons from BPAN cases, one successfully recapitulated the iron accumulation seen in patient brains and further revealed excessive oxidative stress and lysosomal dysfunction, showing that some disease phenotypes may be intrinsic to WDR45-deficient neurons^47^. However, whether neuropathology is impacted by non-cell autonomous factors, and whether this impact is neuroprotective or neurotoxic in BPAN, has not yet been investigated.

In this study, we employed iPSCs derived from BPAN patients and healthy control subjects to investigate the cell autonomous consequences of loss of WDR45 function in microglia. Intriguingly, state-of-the-art approaches including high-performance secretome enrichment with click sugars (hiSPECS)^48^ and NanoString transcriptomic profiling revealed that BPAN patient-derived microglia undergo a profound shift from homeostasis toward chronic activation, marked by dysregulation of immune, stress, and autophagy pathways, impaired lysosomal function, and enhanced antigen presentation. Together, these data suggest that microglial dysfunction may be involved in BPAN pathogenesis by sustaining a maladaptive neuroinflammatory environment, and should be further investigated as a potential component of the disease mechanism.

Our study presents the first patient-derived microglia platform for BPAN research and leverages this valuable tool to shed light on potential mechanisms that may contribute to BPAN neuropathology. We investigated both intrinsic changes to microglial physiology resulting from *WDR45* mutation as well as non-cell autonomous mechanisms potentially driving selective vulnerability of dopaminergic neurons in the SN, taking critical steps towards a more holistic view of neuropathology in BPAN.

## Results

### iPSC-derived microglia from BPAN patients and healthy controls exhibit lineage identity and functional competence

To establish a disease-relevant, patient-derived model for studying microglial function in BPAN, we utilized iPSC lines from two individuals carrying heterozygous *WDR45* mutations, c.519+1_519+3delGTG (BPAN 1)^47^ and c.344+2T>G in exon 6 (BPAN 2), alongside iPSC lines from two healthy controls (control 1, control 2) (Supplementary Fig. 1A-E). Of note, BPAN 1 patient carries an additional heterozygous *PLA2G6* variant (c.91G>A), however, its recessive inheritance pattern makes it unlikely to be pathogenic with a functional wild-type allele present. Thus, WDR45 deficiency is considered the primary cause of disease in this patient.

All four iPSC lines used in this study were thoroughly characterized for pluripotency prior to differentiation into microglia-like cells. Immunocytochemistry demonstrated robust expression of the canonical pluripotency markers SSEA and OCT4 (Supplementary Fig. 1A), as well as SOX2 and NANOG (Supplementary Fig. 1B). Quantitative PCR analysis further confirmed high and comparable expression of key pluripotency genes, including *SOX2* (Supplementary Fig. 1C), *NANOG* (Supplementary Fig. 1D), and *TGDF* (Supplementary Fig. 1E) across all lines, collectively confirming that all iPSC lines maintained a pluripotent state.

iPSCs from BPAN patients and healthy controls were differentiated into microglia-like cells (iMG)^49,50^ (Figure 1A), exhibiting robust expression of canonical microglial markers, as confirmed by RT-qPCR analysis for *TMEM119*, and *P2RY12* (Supplementary Figure 2A-B) or immunoblot analysis for IBA1 (Figure 1B). These classical microglial markers were not detected in iPSC-derived dopaminergic neurons, which instead expressed the neuronal marker tyrosine hydroxylase (TH), absent in iMG of all four lines (Supplementary Figure 2C).

**Figure 1.**
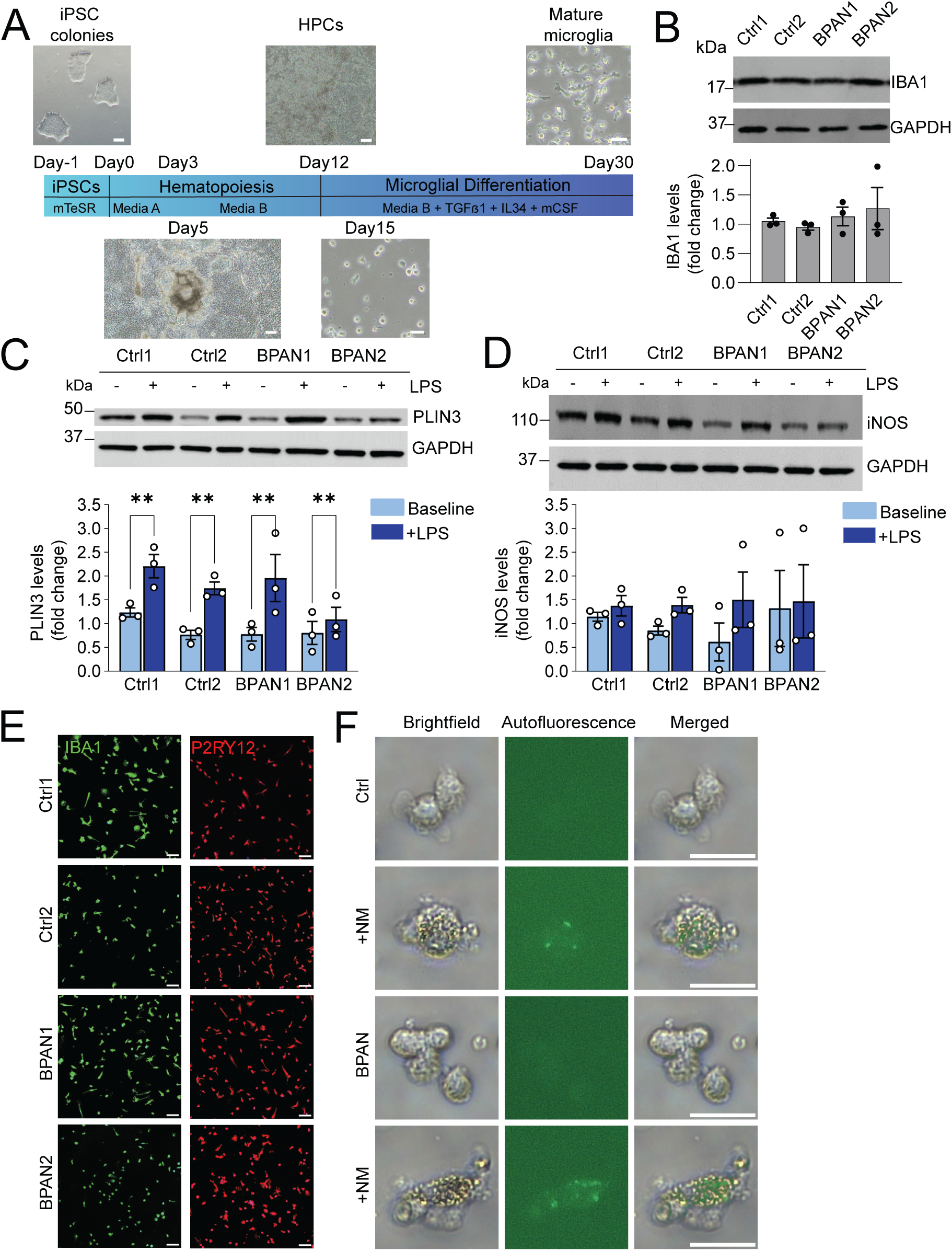
Differentiation and characterization of iPSC-derived microglia. (A) Schematic outline of the microglial differentiation protocol, with representative phase-contrast images of cultures at day 0, day 5, day 12, day 15, and day 30, illustrating key morphological transitions during maturation. (B) Western blot analysis of IBA1 protein expression across all iMG lines (n = 3), with GAPDH as a loading control. Quantification is shown as bar plots (mean ± SEM). (C) Western blot analysis of PLIN3 protein levels in all lines, demonstrating a significant increase following LPS treatment (100 ng/mL, 24 hours) compared to untreated controls. Statistical significance was determined by two-way ANOVA (n = 3) (D) Western blot analysis of iNOS protein levels in all lines after LPS treatment (100 ng/mL, 24 hours), showing no significant change compared to untreated controls. (E) Representative immunofluorescence images of all lines stained for IBA1 (green) and P2RY12 (red). Scale bar, 50 μm. (F) Representative images of control and BPAN patient iMG following neuromelanin treatment (5 ug/mL, 24 hours), showing brightfield, neuromelanin autofluorescence, and merged channels. Scale bar, 10 μm. HPCs, Hematopoietic progenitor cells; NM, neuromelanin; LPS, lipopolysaccharide.

To assess the functional responsiveness of BPAN patient-derived iMG, we evaluated their ability to mount an innate immune response following inflammatory stimulation. PLIN3, a lipid-droplet surface protein marking lipid droplet-accumulating, pro-inflammatory microglia, and iNOS, an enzyme producing nitric oxide during innate immune activation, were selected as complementary indicators of inflammatory responsiveness^51,52^. Immunoblot analyses demonstrated that treatment with lipopolysaccharide (LPS), a bacterial endotoxin that activates innate immune responses, led to significant increase in PLIN3 protein levels (Figure 1C) and persistent expression of iNOS (Figure 1D), suggesting preserved inflammatory responsiveness. In addition, immunocytochemical analysis confirmed that iMG from both control and BPAN patient lines robustly expressed IBA1, a canonical marker of microglial identity (Figure 1E, left panels in green). In parallel, P2RY12, a marker associated with homeostatic and non-activated microglia, was also strongly expressed in all lines at baseline (Figure 1E, right panels in red). The co-expression of IBA1 and P2RY12 validates the successful differentiation of iPSCs into iMG with a homeostatic phenotype.

To further interrogate microglial functionality in disease-relevant conditions, phagocytic competence was assessed by exposing iMG to neuromelanin isolated from human brain^53^. Neuromelanin is a physiological polymer composed of lipids, metals, and proteins that accumulates in nigral dopaminergic neurons during healthy aging, and to an even greater extent in PD. Upon neuron death, neuromelanin is released into the extracellular space, then triggering activation of microglia. Brightfield and fluorescence microscopy demonstrated clear engulfment of exogenously provided neuromelanin, supporting the preservation of phagocytic function in both control and BPAN patient iMG (Figure 1F). In parallel, NLRP3 inflammasome activation was evaluated by immunostaining. Under basal conditions, NLRP3 signal was low, however, following neuromelanin exposure, an increase in NLRP3 signal and the appearance of punctate structures were observed, a classic sign of inflammasome assembly (Supplementary Figure 2E).

Together, these results demonstrate that differentiated iMG from both BPAN patients and controls exhibit key transcriptional, morphological, and functional hallmarks of mature microglia, providing a reliable and disease-relevant model system for downstream mechanistic investigations in BPAN.

### Transcriptomic profiling identifies a disease-associated immune signature in BPAN patient microglia

Transcriptomic profiling offers powerful insights into disease-associated molecular changes. In BPAN, the transcriptomic changes underlying neurodegeneration remain poorly understood. To address this gap, we performed targeted transcriptomic analysis on iMG from BPAN patients and healthy control lines using the NanoString platform, which enables the sensitive detection of immune, apoptotic, and neuroinflammation-related gene expression changes.

Targeted transcriptomic profiling using the Neuroinflammation gene panel revealed significant alterations in gene expression related to key microglial pathways in BPAN patient iMG compared to controls. Heatmap visualization highlighted widespread changes across key microglial pathways including innate and adaptive immunity, cytokine signaling, stress response, microglial homeostasis, and apoptosis, indicating a clear shift from a homeostatic to a reactive, disease-associated state (Fig. 2A). Significant differences in BPAN patient iMG were identified through pathway score analysis using the NanoString Advanced Analysis module, which summarizes the activity of predefined biological pathways based on coordinated expression patterns across grouped target genes, rather than relying solely on individual gene-level changes. Core microglial programs were disrupted, with downregulation of genes associated with microglial homeostasis and surveillance (Fig. 2B), and upregulation of pathways involved in NF-kB signaling, cytokine production, and inflammatory activation (Fig. 2C–E). Notably, the downregulation of autophagy-related genes (Fig. 2F) is consistent with the known role of WDR45 in autophagic regulation.

**Figure 2.**
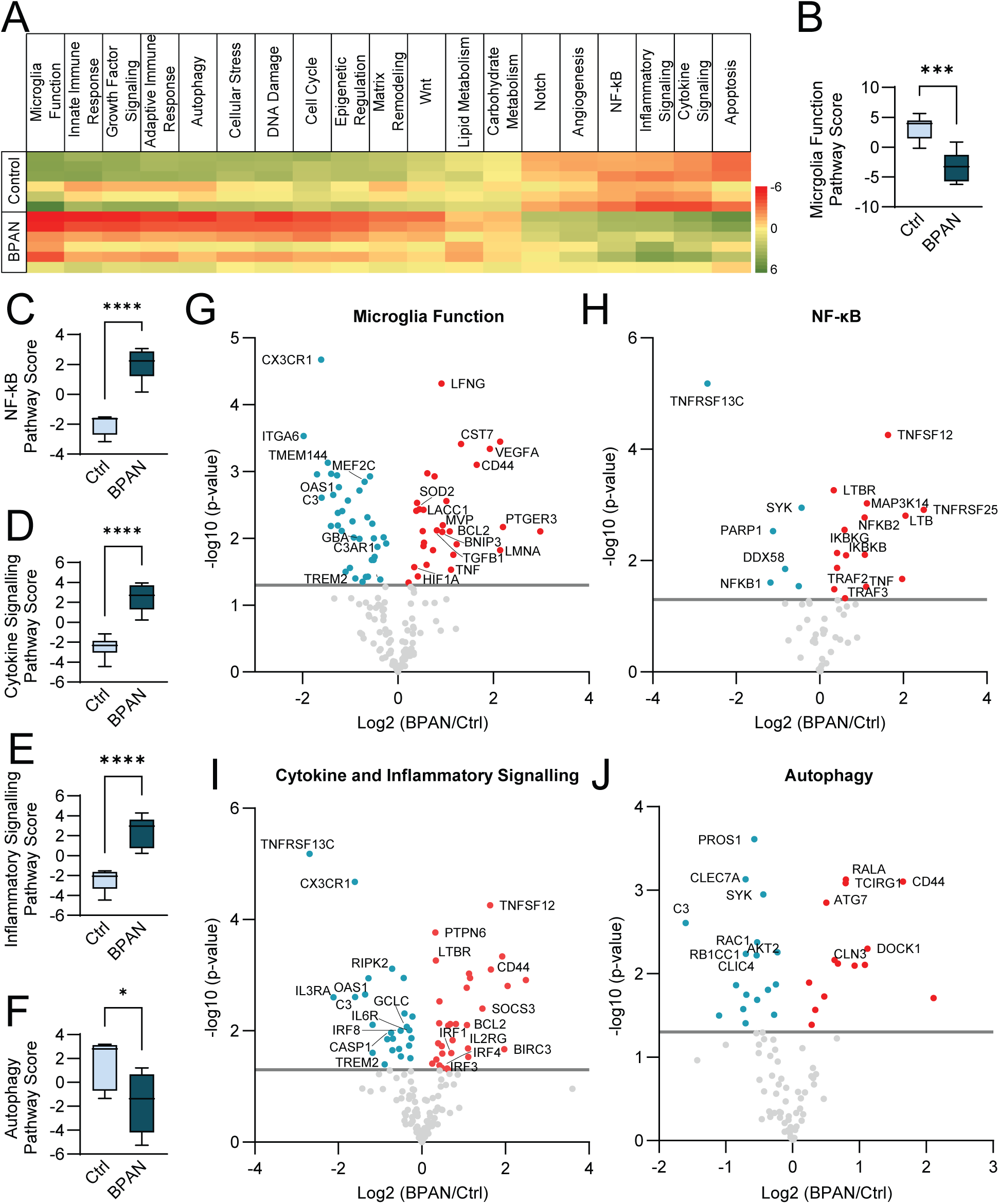
NanoString-based neuroinflammatory pathway analysis shows proinflammatory phenotype. (A) Heatmap displaying pathway scores for 19 relevant neuroinflammation-associated pathways, derived from the NanoString Neuroinflammation Panel, across individual samples (three replicates per line; two control and two BPAN patient lines). Red indicates pathway downregulation and green indicates upregulation relative to baseline. (n = 6 per group, combining replicates from two control and two BPAN patient lines). (**B–F**) Box plots showing pathway scores for selected functional categories, including microglia function (B), NF-κB signaling (C), cytokine signaling (D), inflammatory signaling (E), and autophagy (F). Each box plot displays the distribution of pathway scores (min to max) for all samples in each group (n = 6 per group, combining replicates from two control and two BPAN patient lines). Statistical significance was determined by unpaired t-test; bars represent median and interquartile range. (**G–J**) Volcano plots illustrating differential gene expression within the most relevant pathways: microglia function (G), NF-κB signaling (H), combined cytokine and inflammatory signaling (I), and autophagy (J). Red points indicate upregulated genes, blue points indicate downregulated genes, and light gray points represent nonsignificant changes. The gray horizontal line marks the significance threshold with Benjamini-Yekutieli adjusted p-values (p = 0.05, –log₁₀ = 1.3). Sig, significance.

To further dissect these pathway-level changes, we focused on specific genes within these dysregulated networks. Core microglial maintenance genes (*CX3CR1*, *MEF2C*) were downregulated, while genes linked to inflammation and stress responses, such as *CST7, VEGFA, CD44, PTGER3, TNF*, and *LMNA* were upregulated (Fig. 2G; Supplementary figure 3). This pattern recapitulates a partial disease-associated microglia (DAM) phenotype observed in states of neurodegeneration.

Further, genes involved in microglial homeostasis, immune resolution, and phagocytosis (*ITGA6, OAS1, GBA, C3AR1*) were diminished, alongside widespread downregulation of ribosomal genes, indicating impaired protein synthesis and a metabolically compromised state. Simultaneously, stress adaptation and survival genes (*SOD2, LACC1, MVP, TGFB1, BCL2, BNIP3, HIF1A*) were elevated, consistent with chronic cellular stress. The NF-κB pathway showed broad activation, with upregulation of key pro-inflammatory ligands (*TNF, TNFSF12, LTB*), receptors (*LTBR, TNFRSF25*), and intracellular mediators (*TRAF2, TRAF3, MAP3K14, IKBKB*) (Fig. 2H). Notably, *NFKB1* (*p50*) was diminished, potentially removing critical anti-inflammatory feedback.

Microglial signaling was further altered by suppression of key regulators of microglial activation and immune homeostasis genes (*TREM2, C3, OAS1, IL3RA, IRF8*) and elevation of inflammatory and antigen-presentation markers (*CD44, VEGFA, SOCS3*) (Fig. 2I). The observed alterations to cytokine receptor profiles, including increased *IL2RG*, and decreased *IL6R* and *IL3RA*, suggests altered responsiveness to interleukin signaling. Such shifts in interleukin receptor expression are complemented by changes in interferon regulatory factors (IRFs), which showed an expression pattern aligning with persistent inflammatory signaling, reflecting remodeling of interferon pathways that could prolong or amplify neuroinflammation. A shift toward apoptosis resistance and chronicity rather than acute inflammatory cell death was indicated by the downregulation of key executioner caspases (*CASP3, CASP7*) and inflammatory caspase *CASP1*, alongside upregulation of anti-apoptotic genes (*BCL2, BIRC3*). This pattern was further supported by increased expression of survival-associated regulators such as *MDM2* and *BNIP3*, and reduced levels of pro-apoptotic mediators including *CASP6* and *BIRC2,* which was accompanied by reduced *GCLC* suggesting diminished oxidative stress resilience (Supplementary Figure 3G). DNA damage response pathways showed broad dysregulation, with downregulation of homologous recombination genes such as *RAD51, RAD51C*, and *BARD1*, along with checkpoint mediators *RAD17* and *H2AFX*, suggesting impaired double-strand break repair. While upstream sensors *ATR* and *ATM* were modestly upregulated, downstream repair signaling appeared insufficiently activated indicating compromised genome maintenance capacity, potentially favoring genomic instability and persistent cellular stress (Supplementary Figure 3H).

Autophagy-related genes displayed a mixed regulatory pattern, with compensatory upregulation of *ATG7* contrasted by downregulation of key mediators of autophagy progression (*RB1CC1, CLIC4, RAB7A*) and mitophagy (*PINK1, OPTN*), indicating potential impairment in autophagosome maturation and lysosomal degradation. Similarly, reduced expression of *CLEC7A, SYK* and *PROS1* points to disrupted crosstalk between innate immune signaling and autophagy, which may compromise microglial phagocytic capacity and hinder effective clearance of cellular debris. (Fig. 2J).

In summary, BPAN patient iMG exhibit a deeply altered transcriptomic signature that reflects a shift from neuroprotective to potentially pathogenic behavior. Their chronic activation, impaired resolution mechanisms, and resistance to apoptosis create a persistent neuroinflammatory state.

### Secretome analysis reveals lysosomal dysfunction, altered antigen presentation, and chronic activation of BPAN patient microglia

To assess the paracrine signaling of BPAN patient iMG, we employed hiSPECS, a sensitive mass spectrometry-based approach that enables selective profiling of secreted proteins^48^. This method provides a comprehensive and unbiased snapshot of the secretory activity of living cells under physiological conditions, capturing even low abundance secreted factors (Fig. 3A). Secretome profiling of BPAN patient iMG revealed several converging alterations with potential functional relevance. Most prominently, proteins involved in lysosomal degradation and phagocytic processes were broadly less abundant. This included acid hydrolases such as ARSA, and IDUA (Fig. 3A). This pattern indicates a reduced capacity for degrading phagocytosed material and clearing cellular debris, suggesting compromised microglial housekeeping functions.

**Figure 3.**
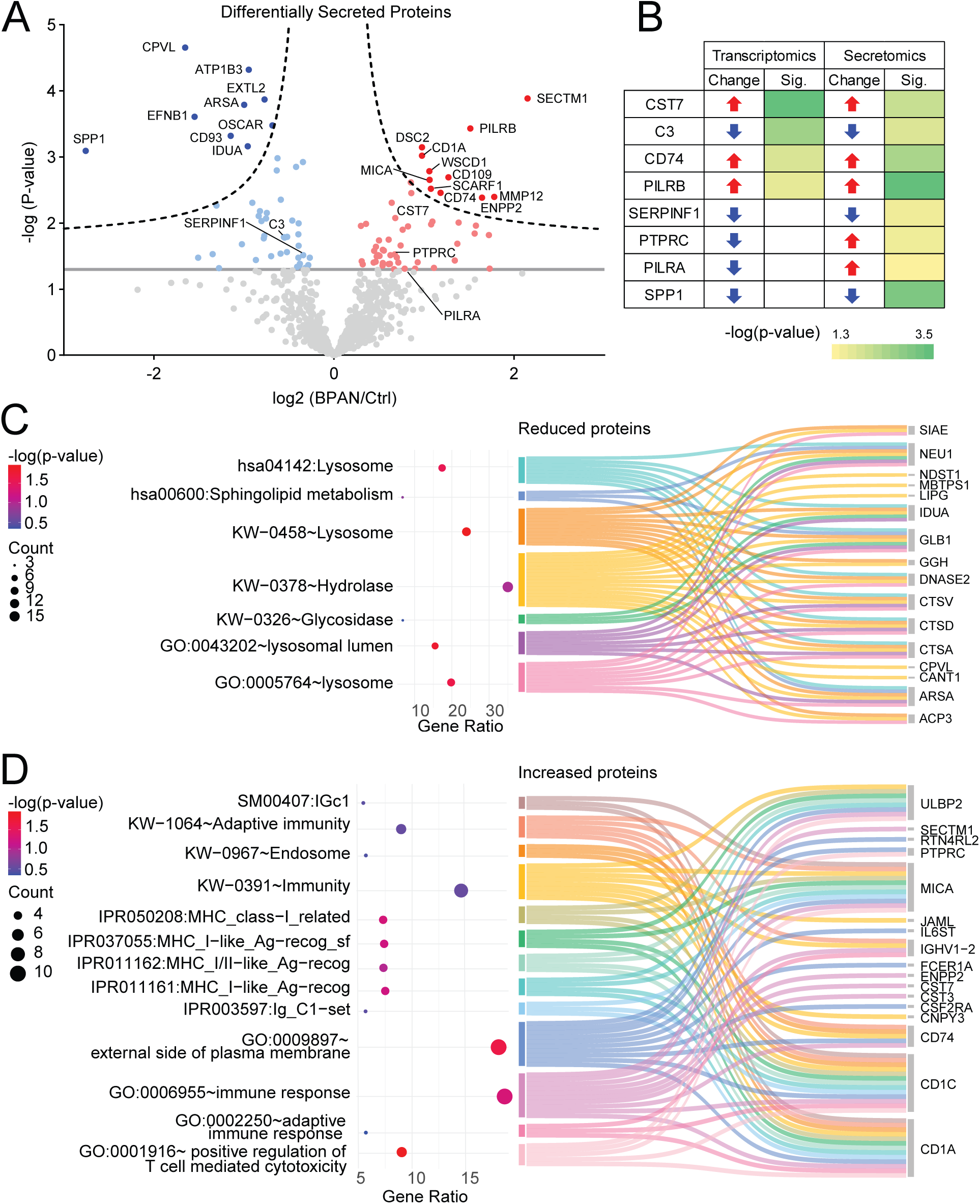
Secretomics analysis of BPAN patient iPSC-derived microglia. (A) Volcano plot of BPAN patient versus control iMG. The -log10 transformed p-values of each protein are plotted against their log2 fold change transformed protein label free quantification ratios. Proteins significantly more secreted from BPAN are displayed as red filled dots, while less secreted proteins are displayed as blue filled dots. Unaltered proteins are displayed as the gray dots. Proteins are labeled with their UniProt gene names. (B) Comparison of the differentially secreted proteins to the targeted transcriptomics. The increase in gene expression and secretion is represented with green upper arrow, decrease with lower red arrow and p-values as color gradient. (C) Group of terms in cluster 2 of Pathway enrichment analysis for downregulated proteins. The dot plots (left) show the total gene number of a term in percentage as dot size and the p values as a color gradient. Sankey plots (right) show how each term relates to each gene. (D) Group of terms in cluster 5 of Pathway enrichment analysis for upregulated proteins. The dot plots (left) show the total gene number of a term in percentage as dot size and the p-values as a color gradient. Sankey plots (right) show how each term relates to each gene.

In parallel, there was a marked reduction of the DAM marker osteopontin (SPP1), which is typically elevated in other neurodegenerative conditions and has been linked to neuroprotective and reparative functions^54^. This may reflect a dysfunctional or incomplete transition of BPAN patient iMG into neuroprotective or reparative DAM phenotypes.

Conversely, several proteins involved in immune activation and antigen presentation showed a higher abundance in BPAN patient iMG in secretome analysis (Fig. 3A). These included the transmembrane proteins SECTM1, CD1A, CD74, and MICA, which are released by proteolytic ectodomain shedding, suggesting a shift toward a more pro-inflammatory and antigen-presenting phenotype. In addition, increased secretion of matrix metalloproteinase MMP12 points to enhanced extracellular matrix remodeling, potentially increased microglial motility or invasiveness, and additional support for an incomplete or atypical DAM phenotype.

Integration of transcriptomic (Fig. 2) and secretomic data revealed a subset of genes, i.e. *CST7*, *C3, CD74*, and *PILRB,* that showed consistent changes at both the mRNA and secreted protein levels, underscoring their likely central role in BPAN microglial dysfunction (Fig. 3B). Other proteins, such as SERPINF1, PTPRC (CD45), PILRA, and SPP1, were only differentially abundant at the secretomics level indicating alterations in protein secretion. Both SERPINF1 and SPP1 trended in the same direction at the transcript level, though not reaching statistical significance. Notably, the strong reduction in secreted SPP1, despite stable transcript levels, suggests additional regulation through translational repression, increased protein instability or degradation, or impaired secretion.

Follow-up pathway enrichment analysis of secreted proteins from BPAN patient and control iMG revealed several alterations to functionally distinct clusters (Fig. 3C, D). Cluster 2, comprising proteins reduced in BPAN patient microglial secretome, was enriched for lysosomal components and enzymatic activities essential for macromolecule degradation, including glycosidases and hydrolases (Fig. 3C). Key lysosomal hydrolases such as CTSD, GLB1, IDUA, and DNASE2 were among the proteins showing reduced secretion. The concurrent enrichment of sphingolipid metabolism pathways, involving enzymes like ARSA, further suggests disrupted lipid turnover. These alterations indicate impaired lysosomal function and extracellular degradative capacity, consistent with compromised autophagic clearance and defective processing of phagocytosed material, key processes for microglial maintenance of neural tissue homeostasis. Proteins with higher abundance in BPAN patient microglial secretomes (Cluster 5) were strongly enriched for immune-related pathways, including adaptive immune response, positive regulation of T cell mediated cytotoxicity, and broader immune response processes (Fig. 3D). For example, CD74 and CD1C, key molecules involved in antigen processing and presentation, reflect enhanced capacity for T cell activation. Correspondingly, cytokine receptor subunits such as CSF2RA and IL6ST highlight increased microglial responsiveness to inflammatory signals. The upregulation of immune surveillance ligands MICA and ULBP2, protease inhibitors CST3 and CST7, immune modulators SECTM1 and JAML, along with receptors such as FCER1A and PTPRC (CD45), collectively reflects enhanced regulation of extracellular proteolysis, immune signaling, and cell-cell interactions in this cluster.

Together, these findings define a distinct secretory profile in BPAN patient iMG marked by impaired lysosomal and phagocytic function, loss of neuroprotective DAM features, upregulation of antigen presentation, and chronic immune activation, potentially contributing to neuroinflammatory pathology.

## Discussion

Our multiomics analysis of BPAN patient-derived iMG reveals a complex and divergent activation profile that only partially overlaps with the DAM phenotype, thereby expanding BPAN research beyond neurons and autophagy to highlight the importance of glial contributions and neuroimmune mechanisms. Notably, this profile includes downregulation of homeostatic genes such as *CX3CR1* and *MEF2C*, crucial for synaptic pruning and neuron–microglia communication^55,56^. In parallel, upregulation of inflammatory and remodeling genes like *CD44, VEGFA*, and *C3* indicates a shift toward a reactive, neuroinflammatory phenotype^57,58^. In addition, downregulation of *IRF8*, disrupts homeostatic maintenance, phagocytic function, and inflammatory regulation^59,60^. The result is a shift toward states that may impair neuronal function or exacerbate neurodegeneration.

Both canonical and non-canonical NF-κB signalling are activated, reflected by upregulation of *MAP3K14* and IKBKB, and ligands such as *TWEAK (TNFSF12)* and *LTB* and its receptor, which are known drivers of microglial activation in diseases like multiple sclerosis^61,62^.

Downregulation of antiviral genes like *OAS1, RSAD2*, and *DDX58* suggests impaired innate immune defense, likely reducing their ability to detect and respond to viral pathogens with type I interferon signaling^63^. Upregulation of survival genes such as *BCL2* and downregulation of *PARP1* suggest apoptosis resistance and senescence^64^. Functionally, this may lead to the persistence of dysfunctional, pro-inflammatory microglia in the brain, contributing to chronic neuroinflammation.

Lastly, altered expression of autophagy-related genes (*ATG7, CLEC7A, PROS1*) may reflect compensation for WDR45-associated autophagy dysfunction, supporting defective clearance mechanisms may contribute to progression of neurodegeneration in BPAN^34,42^.

To further explore the functional consequences of the transcriptional changes observed in BPAN patient iMG, we integrated these findings with secretomic data. This combined approach allowed us to assess whether dysregulated gene expression translated into altered protein secretion, which is critical for understanding microglial communication and effector function. BPAN patient iMG displayed a distinct secretory signature marked by impaired proteostasis, altered immune signaling, and chronic activation.

The coordinated low abundance secretion of lysosomal enzymes, including CTSD, GLB1, IDUA, and NEU1, alongside the suppression of the complement component C3, indicates impaired microglial clearance of apoptotic material and protein aggregates, a hallmark of microglial dysfunction in neurodegenerative diseases^65,66^ and a process critical for maintaining CNS homeostasis and preventing sterile inflammation^1,40,67^. These lysosomal deficits likely reflect a disruption in autophagy-linked exocytosis, a cellular process previously shown to be impaired in WDR45-deficient non-microglial cells, and here demonstrated for the first time in patient-derived microglia^34,40,47^. In parallel, the strong reduction of secreted SPP1, a protein consistently upregulated in DAM profile, suggests an atypical or incomplete DAM phenotype^54^. This blunted DAM-like response may reflect a failure to mount neuroprotective programs, further implicating lysosomal dysfunction as a key driver of compromised microglial activity. The increased secretion of immune mediators, including antigen presentation molecules CD74, MICA, CD1C, and ULBP2, as well as cytokine receptor CSF2RA and protease MMP12, points to a reactive microglial state characterized by enhanced immune signaling, antigen processing, and extracellular matrix remodeling^68,69^. This pattern suggests a shift toward a pro-inflammatory, antigen-presenting phenotype that may sustain or amplify chronic neuroinflammatory signaling in BPAN. Notably, this profile parallels microglial phenotypes described in neurodegenerative disorders such as PD and AD, where persistent microglial activation exacerbates neuronal injury^53,70,71^. These changes may represent either a compensatory attempt to respond to cellular damage or a maladaptive immune overactivation driven by lysosomal stress. When considered alongside the impaired clearance of apoptotic material and protein aggregates, this dual signature of dysfunction, combining defective degradative capacity with heightened immune alertness, provides compelling evidence for a direct contribution of microglia to BPAN pathology. These findings further support emerging models positioning dysfunctional microglia as central players in neurodegeneration, reinforcing the rationale for therapeutic strategies aimed at restoring microglial homeostasis and function.

The overlap between transcriptomic and secretomic changes, such as those involving *CST7, CD74*, and *PILRB*, reinforces the functional relevance of these immune and lysosomal pathways and highlights their central role in BPAN microglial pathology. In contrast, the discordance observed in genes like *SPP1* and *PILRA* underscores the complexity of disease regulation and the importance of post-transcriptional mechanisms in shaping the microglial phenotype in BPAN^72,73^.

BPAN patient-derived iMG reveal a partial DAM phenotype marked by selective activation of certain DAM-associated pathways and suppression of others. Key regulators such as *TREM2* and *CLEC7A* were significantly downregulated, as were homeostatic markers *CX3CR1* and *MEF2C*, indicating disruption of canonical DAM initiation and loss of resting microglial identity. In contrast, inflammatory and lysosomal genes including *ITGAX, CST7*, and *CD68* were upregulated, reflecting a shift toward immune activation. Several other canonical DAM genes (*APOE, LPL, GPNMB, AXL, TYROBP, CSF1R*) remained unchanged, underscoring the incomplete nature of this transition. Overall, this expression pattern suggests a dysregulated, context-specific DAM program in BPAN patient iMG, combining elements of inflammatory activation with impaired lipid- and phagocytosis-related functions.

While the present study provides the first-of-its-kind transcriptomic and secretomic characterization of BPAN patient-derived iMG, several limitations should be acknowledged. First, iPSC-derived microglia are powerful tools, but may not fully capture the complexity of microglial development and function within the 3D CNS environment, including interactions with neurons, astrocytes, vasculature, and the extracellular milieu. Second, the cross-sectional nature of our data does not address dynamic changes over time or the bidirectional nature of microglial communication that shape their phenotype and function *in vivo*. Finally, while transcriptomic and secretomic profiles provide insight into altered pathways, further functional validation will be required to establish causal links between specific dysregulated programs and disease progression. Addressing these limitations in future studies, for example, by incorporating co-culture or organoid models, performing functional assays, correlating findings with *in vivo* models, will be important to more fully define the contribution of microglia to BPAN pathogenesis.

To date, BPAN research has focused predominantly on neuronal and autophagy-related mechanisms, with relatively little attention paid to the role of glia or the neuroimmune system. By shifting focus to patient-derived microglia, our study begins to fill this gap, demonstrating that BPAN patient iMG exhibit a dysregulated molecular signature. Overall, our study emphasizes that BPAN patient iMG are locked in a chronic maladaptive inflammatory state and should be considered as likely contributors to neurodegeneration and promising therapeutic targets in future investigations.

## Materials and Methods

### Subjects

Included in this study are iPSCs from two BPAN patients (BPAN 1 = L-8172^47^; BPAN 2 = 119129, registered at https://biobanknetwork.telethon.it/Sample/View?sampleId =119129) and two healthy control individuals (control 1 = SFC156-03-012, registered at hPSCreg: https://hpscreg.eu/cell-line/STBCi101-A; control 2= SFC065-03-05, registered at hPSCreg: https://hpscreg.eu/cell-line/STBCi057-B). The study was approved by the Ethics Committee of the Medical Faculty of LMU Munich, Germany, and all participants gave written informed consent.

### Culture and characterization of human iPSCs

iPSC cultures were grown as colonies and maintained on Cultrex-coated (R&D Systems) plates and antibiotic-free mTESR Plus medium (Stem Cell Technologies) was refreshed every other day. Cells were manually passaged every 5-7 days.

BPAN and control iPSC lines were characterized for expression of pluripotency markers by quantitative RT-PCR (SOX2, NANOG, TDGF1). RNA was extracted from subconfluent iPSC colonies using the RNeasy kit (Qiagen), then reverse transcribed into cDNA using the iScript ™ cDNA Synthesis Kit (Biorad). Quantitative RT-PCR was performed with SYBR GreenER (Thermo Scientific) on the 7500 Fast Real-Time PCR system (Applied Biosystems). Primer sequences were designed with Primerblast and purchased from Integrated DNA Technologies (Table 1). Ribosomal protein L13 (RPL13) was used as housekeeping gene. All lines were routinely tested for mycoplasma contamination.

**Table 1:**
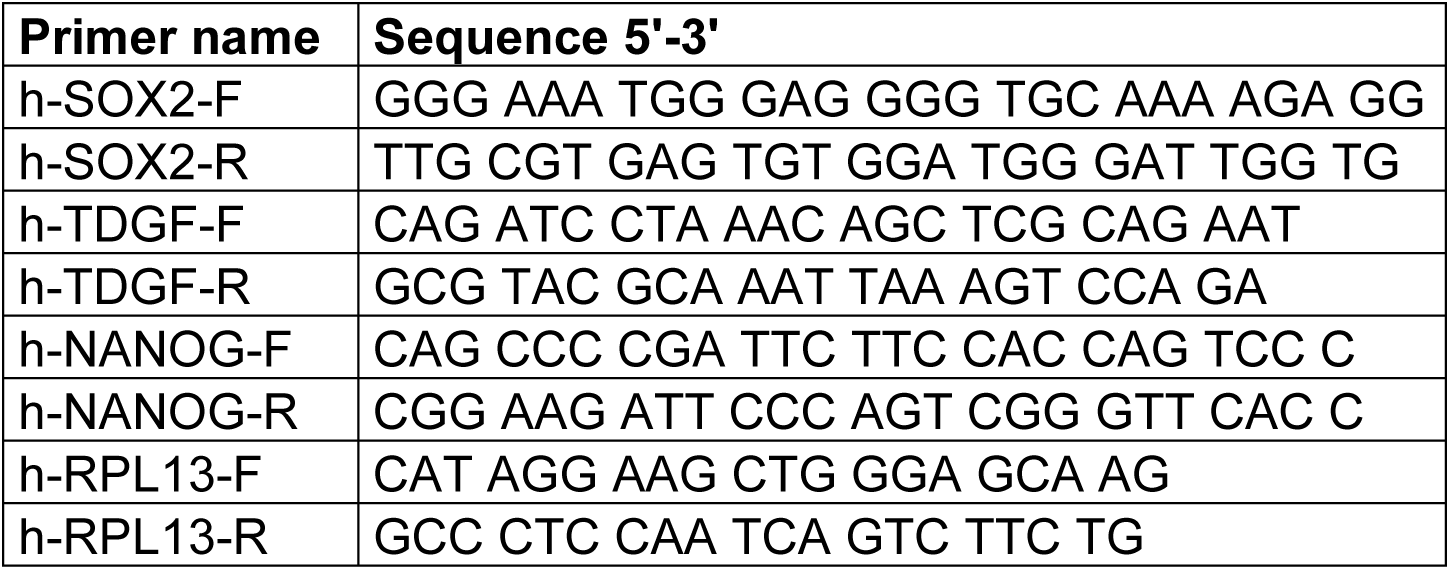
List of RT-qPCR primers used for pluripotency validation of iPSC lines.

### Differentiation and characterization of human iPSC-derived microglia culture

iPSC-derived microglia (iMG) were generated from BPAN patients and controls according to the protocol by McQuade et al., 2018 with modifications^49,50^. Briefly, 4-6 medium-sized iPSC colonies were manually seeded onto a Cultrex-coated six-well plate with fresh mTeSR™ Plus medium. After 18-24 hours, wells containing approximately 4-10 attached colonies/cm² were used for hematopoietic differentiation. Hematopoietic progenitor cells (HPCs) were generated using the STEMdiff hematopoietic kit (STEMCELL Technologies) following the manufacturer’s protocol for 12 days. On day 12, HPCs were gently harvested and seeded into 6-well plates at a density of 1-2×10^5 cells per well in iMG differentiation medium (phenol-free DMEM/F12 supplemented with 2X B27 (Thermo Fisher Scientific), 2X insulin-transferrin-selenite (Thermo Scientific), 0.5X N2 (Life Technologies), 1X GlutaMAX (Life Technologies), 1X MEM non-essential amino acids solution (Thermo Scientific), 5 μg/mL human insulin (Sigma Aldrich), and 400μM α-thioglycerol (Sigma Aldrich). The microglia differentiation medium was supplemented with the following factors for microglia lineage commitment: IL-34 (100 ng/ml; Peprotech), TGF-β1 (50 ng/ml; Peprotech), and M-CSF (25 ng/ml; Peprotech). iMG were maintained in this media until day 30, when the cells were considered mature.

BPAN patient and control iMG were characterized by quantitative RT-PCR (TMEM119, P2RY12). RNA extraction was performed using the RNeasy kit (Qiagen) which was then reverse transcribed into cDNA with the iScript ™ cDNA Synthesis Kit (Biorad). Quantitative RT-PCR was performed with SYBR GreenER (Thermo Scientific) on the 7500 Fast Real-Time PCR system (Applied Biosystems). Primer sequences were designed with Primerblast and purchased from Integrated DNA Technologies (Table 2). ALU repeat units was used as housekeeping gene.

**Table 2:**
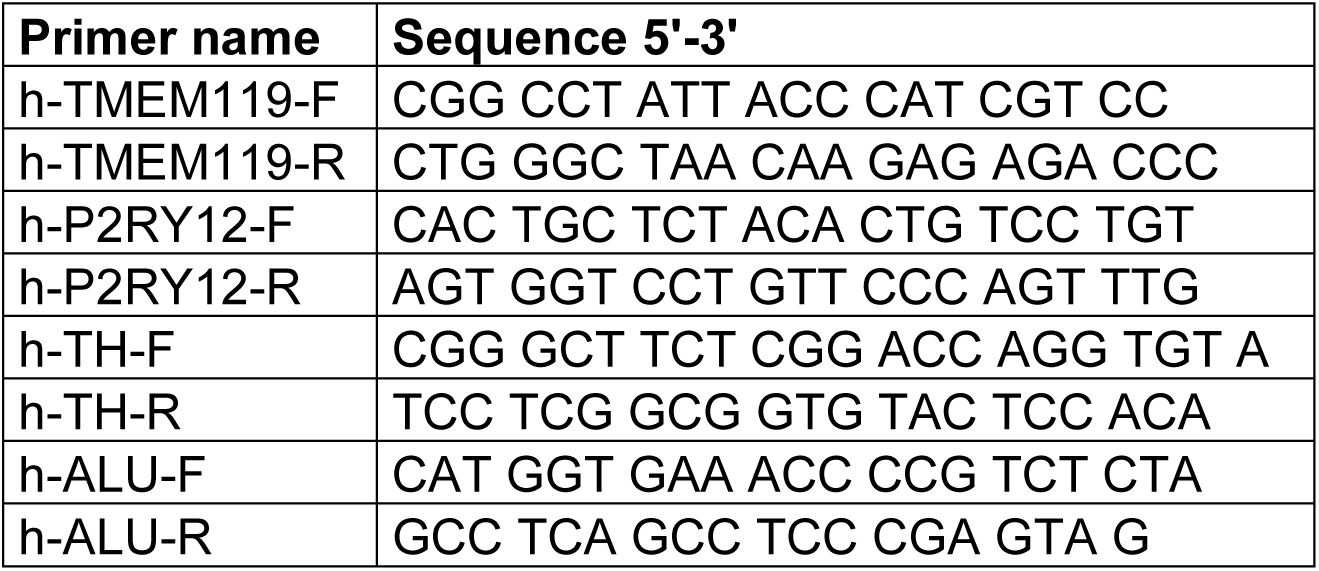
List of RT-qPCR primers used for validation and characterization of iPSC-derived microglia.

### Differentiation of human iPSCs into midbrain dopaminergic neurons

The differentiation of midbrain dopaminergic neurons was done as previously described with minor adaptations^74^. In detail, healthy control human iPSCs were cultured on Cultrex-coated plates in antibiotic-free mTeSR Plus medium (StemCell Technologies) and manually passaged every 5–7 days. Prior to passaging, spontaneously differentiated regions were removed under a stereomicroscope, and colonies were mechanically fragmented using a pipette tip. Aggregates were transferred to fresh Cultrex-coated wells and maintained at 37°C with 5% CO₂, with medium changes every 48 hours. Neuron differentiation was initiated when cultures reached over 95% confluence (Day 0). On day 0, the medium was changed to KSR differentiation medium composed of KnockOut DMEM (Thermo Fisher), 15% KnockOut Serum Replacement (Thermo Fisher), 1× GlutaMAX (Thermo Fisher), 1× penicillin-streptomycin (Thermo Fisher), and 1 mM β-mercaptoethanol (Sigma-Aldrich), supplemented with dual SMAD inhibitors LDN193189 (LDN) and SB431542 (SB). On day 1, Sonic hedgehog protein (SHH), purmorphamine, and Fibroblast Growth Factor 8 (FGF8) were added, followed by CHIR99021 (CHIR) on day 3 to promote floor plate patterning. On day 5, cells were transitioned to NbSm1 medium, consisting of Neurobasal Medium (Thermo Fisher), NeuroCult SM1 supplement (STEMCELL Technologies), 1× GlutaMAX (Thermo Fisher), and 1× penicillin-streptomycin (Thermo Fisher), supplemented with LDN193189, SHH, purmorphamine, FGF8, and CHIR. From day 7, SHH, purmorphamine, and FGF8 were withdrawn, and only LDN and CHIR were maintained. Between days 11–14, cells were passaged as 1–2 mm chunks onto poly-D-lysine/laminin-coated dishes in NbSm1 medium supplemented with CHIR, BDNF, ascorbic acid, GDNF, TGF-β3, dibutyryl-cAMP, and DAPT. Between days 25–30, cells were dissociated into single-cell suspensions using Accutase and replated on coated dishes in the same media formulation. From day 40 onward, cells were maintained in NbSm1 medium without supplements until collection.

### Immunofluorescent staining and image analysis

For immunocytochemical analysis, all iPSCs lines were seeded onto Cultrex-coated (R&D Systems) glass coverslips and iMG were seeded onto uncoated glass coverslips, fixed in 4% paraformaldehyde and permeabilized with 0.3% Triton X-100 in PBS. Cells were then blocked for 1 hour at room temperature in a solution containing 2% BSA and 5% normal goat serum in PBS-Triton. Coverslips were then incubated overnight at 4°C in a humidified chamber with primary antibodies.

iPSCs were characterized for pluripotency status via immunocytochemical analysis using the following primary antibodies: anti-Oct4 (Abcam #ab19857, 1:100), anti-SOX2 (Abcam #ab79351, 1:100), anti-Nanog (Abcam #ab80892, 1:100), anti-SSEA-4 (Millipore #MAB4304, 1:100). iMG were characterized using primary antibodies at the following dilutions: anti-IBA1 (Abcam, #ab178846, 1:500), anti-P2RY12 (Sigma Aldrich, #A014518, 1:200), anti-NLRP3 (Santa Cruz, #sc-518123). After washing with PBS, cells were then incubated with Alexa-conjugated anti-rabbit and Alexa-conjugated anti-mouse secondary antibodies (1:500) for 1 hour at room temperature. Following another wash step, DAPI (Merck, #MBD0015) was applied for 10 minutes at room temperature, coverslips were mounted onto glass microscope slides using ProLong™ glass antifade mounting media (Thermo Fisher), and allowed to dry overnight. Images for iMG were captured keeping the settings constant for each antibody using the confocal microscope (Leica TCS SP5, Leica Microsystems, Wetzlar, Germany). Pictures for iPSCs characterization were taken by fluorescence microscope (ECHO Revolve, Discover Echo, San Diego, CA, USA). At least 3 fields of view were analyzed per coverslip.

### Protein extraction and western blot analysis

Cultured iMG were harvested by collecting both floating and adherent iMG into the same tube. The adherent cells were dissociated by incubation with TrypLE for 5 minutes at 37°C, then all cells were centrifuged at 250 x g for 5 minutes. The resulting pellet was washed 1X with PBS then centrifuged again at 500 x g for 10 minutes for collection. Cell pellets were sequentially homogenized using RIPA lysis buffer (Thermo Scientific) containing protease-phosphatase inhibitor cocktail (Roche). 10ug of each protein sample was run via SDS-PAGE on gradient gels under reducing conditions. Gels were blotted onto polyvinylidene difluoride (PVDF) membranes and blocked with Intercept blocking reagent (Li-Cor) for 1 hour at room temperature. Primary antibodies used for western blotting were: anti-GAPDH (Millipore #MAB374, 1:10,000) anti-IBA1 (Abcam #ab178846, 1:500), anti-PLIN3 (Proteintech #10694-1-AP, 1:500), and anti-iNOS (Proteintech #18985-1-AP, 1:500). Blots were washed 3x with 1X TBS-T, probed with the appropriate fluorescent secondary antibody (IRDye, Li-Cor), washed 3x with 1X TBS-T and imaged with the Odyssey XF Dual-Mode Imaging System (Li-Cor).

### LPS and neuromelanin treatment of iMG

iMG were plated at a density of 50,000 cells/cm2 on uncoated 6-well plates and cover slips. After allowing the cells to attach for 24 hours, they were treated with 100 ng/ml of LPS (Thermo Scientific) for 24 hours. For western blot analysis, both floating and adherent iMG were collected into the same tube. The adherent cells were dissociated by incubation with TrypLE for 5 minutes at 37°C, then all cells were centrifuged at 250 x g for 5 minutes. The resulting pellet was washed 1X with PBS then centrifuged again at 500 x g for 10 minutes for collection.

For treatment with neuromelanin, mature iMG were plated at a density of 50,000 cells/cm² on uncoated 24-well plate coverslips and allowed to attach for 24 hours. Neuromelanin suspension was prepared from a purified human neuromelanin stock (0.5 mg/mL) stored at –80 °C. To sterilize the neuromelanin, approximately 0.5 mL of ethanol was added directly into the glass tube containing the neuromelanin, gently shaken for 5 minutes, and evaporated to dryness under a nitrogen stream. Sterile distilled water was then added to restore the original neuromelanin concentration (0.5 mg/mL). The tube was sealed tightly and sonicated for 5–10 minutes to break up larger granules, with additional manual dispersion using the tip of a glass pipette as needed. The neuromelanin suspension was then gently shaken at room temperature, protected from light, for 3–4 days until fully rehydrated and no floating granules remained. Vortexing or tube inversion was avoided to prevent neuromelanin adherence to the tube walls. Prior to treatment, the suspension was gently mixed, and wide-bore pipette tips (cut to avoid clogging) were used to transfer aliquots. The neuromelanin suspension was pre-incubated in complete culture medium for 2–3 hours to allow interaction with medium components before being applied to the cells at a final concentration of 5 µg/mL for 24 hours. Brightfield images were acquired using a Nikon Eclipse Ts2 inverted microscope equipped with a TrueChrome Metrics digital camera.

### Targeted transcriptomic analysis using the NanoString neuroinflammation panel

Targeted gene expression profiling was performed using the nCounter Human Neuroinflammation Panel v1.0 (NanoString Technologies, Seattle, WA, USA), which includes 770 curated mRNA probes related to key neuroinflammatory pathways.

Total RNA was extracted from frozen iMG pellets using the RNeasy Mini Kit (Qiagen, Hilden, Germany) following the manufacturer’s protocol, and samples were diluted to a final concentration of 20 ng/μL. For each reaction, 100 ng of RNA was hybridized with capture and reporter probes overnight at 65 °C for 16 hours.

Following hybridization, samples were transferred to German Cancer Consortium (DKTK) core facility, where post-hybridization processing and data acquisition were performed using the nCounter Analysis System (NanoString Technologies), in accordance with the manufacturer’s instructions. Raw digital count data were analyzed using nSolver™ Analysis Software (v4.0) and the Advanced Analysis module.

Normalization of mRNA counts was carried out using NanoString’s automated reference gene selection method, which identifies a subset of the most stable housekeeping genes from the panel. Background correction was based on the included negative control probes, and all samples met quality control criteria. Differential expression analysis was conducted using linear modeling implemented in the Advanced Analysis module, with significance thresholds set at a Benjamini– Hochberg adjusted p-value < 0.05.

Pathway-level analysis was performed using the panel’s integrated pathway scoring algorithm, which calculates the first principal component of gene sets associated with each predefined neuroinflammatory pathway to provide an overall pathway activity score per sample.

### Secretomics

Cells were culture in 50 µM N-azido-mannosamine (ManNAz) for 48 hours. The media was collected and filtered with a 0.22 µm spin filter (Costar Spin-X, Sigma Aldrich, CLS8160).

The supernatants were washed two times with 250 µL PBS on 10K Vivacon filters (Sartorius, Germany) and concentrated to 50 µL. Biotinylation of labeled glycoproteins was performed by adding 5 µL 10 mM Sulfo-dibenzocyclooctyne-biotin (Sigma Aldrich, 760706) and incubation for 16 hours at 4°C while shaking at 750 rpm. Afterwards, samples were washed three times with PBS, concentrated to 50 µL and recovered by reverse spinning.

The supernatants were subjected to a lectin-based enrichment using 50 µL concanavalin A (ConA) beads (Merck, C7555) per sample. Glycoproteins were bound to the beads in 1 mL binding buffer (5 mM MgCl_2_, 5 mM MnCl_2_, 5 mM CaCl_2_, 0.5 M NaCl, 20 mM Tris-HCl pH 7.5) rotating end-over-end for at least 2hours at 4°C followed by two washing steps with 1 mL binding buffer. Proteins were eluted in 500 µL elution buffer (500 mM Methyl-α-D-mannopyranoside, 10 mM EDTA, 20 mM Tris-HCl pH 8.5) and Triton X-100 was added to a final concentration of 1% (v/v).

Glycoprotein enrichment was performed with 10 µL of magnetic Streptavidin beads (Thermo Fisher Scientific, 88817) per sample. Beads were washed three times with 1 mL 1% (v/v) Triton-X before they were added to the samples. Then, samples were incubated rotating end-over-end for 2hours at 4°C. The streptavidin beads with the bound glycoproteins were washed three times with SDS buffer (1% (w/v) SDS, 250 mM NaCl, 100 mM Tris-HCl pH 8.5), three times with urea buffer (3M urea, 0.1% sodium deoxycholate (SDC), 50 mM ammonium bicarbonate (ABC)) and once with 0.1% SDC in 50 mM ABC using a Dynamag 2 magnetic rack (Thermo Fisher Scientific, US).

Protein reduction was performed with 100 µL 10 mM dithiothreitol in 0.1% SDC and 50 mM ABC and incubation for 30 minutes at 37°C while shaking at 1000 rpm. The supernatant was discarded and protein alkylation was performed with 100 µL 40 mM iodoacetamide in 0.1% SDC and 50 mM ABC and incubation for 30 min at 24°C in the dark while shaking at 1000 rpm. After removal of the supernatant, on-bead protein digestion was performed in 60 µL 0.1% SDC and 50 mM ABC with 0.2 µg LysC (Promega) for 3 hours at 37°C and subsequently with 0.2 µg trypsin (Promega) for 16 hours at room temperature.

The supernatant was collected and peptides were purified adding 200 µg of a one-to-one mixture hydrophilic and hydrophobic carboxylate coated SeraMag speedbeads (Cytiva, US) using a magnetic rack. Peptide retention on the beads was facilitated by adding acetonitrile (Sigma Aldrich) to a final concentration of more than 95% (v/v) followed by one washing step with 100% (v/v) acetonitrile. After removal of the supernatant, Peptides were eluted in 40 µL 0.1% (v/v) formic acid (FA, Sigma Aldrich) and filtered through 0.22 µm spin filters (Costar Spin-X, Sigma Aldrich, CLS8160). Samples were dried by vacuum centrifugation and dissolved in 20 µL 0.1% formic acid directly before starting the mass spectrometric analysis.

### Mass spectrometry

A volume of 8 µL per sample was separated on a nanoElute nanoHPLC system (Bruker, Germany) on an in-house packed C18 analytical column (15 cm × 75 µm ID, ReproSil-Pur 120 C18-AQ, 1.9 µm, Dr. Maisch GmbH) using a binary gradient of water and acetonitrile (B) containing 0.1% formic acid at flow rate of 300 nL/min (0 min, 2% B; 2 min, 5% B; 62 min, 24% B; 72 min, 35% B; 75 min, 60% B) and a column temperature of 50°C. The nanoHPLC was online coupled to a TimsTOF pro mass spectrometer (Bruker, Germany) with a CaptiveSpray ion source (Bruker, Germany). A Data Independent Acquisition Parallel Accumulation–Serial Fragmentation (diaPASEF) method was used for spectrum acquisition. Ion accumulation and separation using Trapped Ion Mobility Spectrometry (TIMS) was set to a ramp time of 100 ms. One scan cycle included one TIMS full MS scan with 26 windows with a width of 27 m/z covering a m/z range of 350-1001 m/z. Two windows were recorded per PASEF scan. This resulted in a cycle time of 1.4 seconds.

### Mass spectrometry data analysis

The software DIA-NN version 1.8.1 was used to analyze the data ^75^. The raw data was searched against a one protein per gene database from homo sapiens (UniProt, 21758 entries, download: 2024-07-30) and a database with potential contaminants (246 entries) from Maxquant ^76^ using a library free search. Trypsin was defined as protease and two missed cleavages were allowed. Oxidation of methionines and acetylation of protein N-termini were defined as variable modifications, whereas carbamidomethylation of cysteines was defined as fixed modification. Variable modifications were restricted to two per peptide. The precursor and fragment ion m/z ranges were limited from 350 to 1001 and 200 to 1700, respectively. Precursor charge states were set to 2-4 charges. An FDR threshold of 1% was applied for peptide and protein identifications. The mass accuracy for MS1 and MS2 were set to 15 ppm. Ion mobility windows were automatically adjusted by the software. The match between runs and RT-dependent cross-run normalization options were enabled.

The software Perseus version 1.6.2.3 ^77^ was used for further data analysis. Contaminants were removed. Protein label-free quantification (LFQ) intensities were log2 transformed. Changes in protein abundances between the different groups were calculated and a two-sided Student’s Ttest was applied comparing the log2 transformed LFQ intensities between the groups. At least three valid values per group were required for statistical testing. To account for multiple hypotheses, a permutation-based FDR correction ^78^ was applied.

### Pathway enrichment analysis for secretomics

Differentially expressed proteins were defined by a p-value <0.05 for both BPAN patient lines versus both controls. Pathway enrichment analysis was performed using the web-based program DAVID version 6.8 and the Functional Annotation Clustering tool. Gene lists for either increased- or reduced proteins were compared to all proteins relatively quantified in LC-MS/MS in all cell lines, defined as experimental background.

### Statistical analysis

Student’s *t*-test analysis and ANOVA followed by Tukey’s post-hoc test was performed. *P*-values less than 0.05 were considered significant. All errors bars shown in the figures are standard error of the mean (SEM) and asterisks denotes p < 0.05 (*), p < 0.01 (**), p < 0.001 (***) and p < 0.0001 (****).

## Supporting information

Supplemental Table 1: Secretomics Analysis

Supplemental Table 2: NanoString Analysis

## Acknowledgment

This work was supported by a postdoctoral scholarship by the Alexander von Humboldt Foundation (to R.M.W.), by the NBIA Disorders Association (NBIADA, USA), Associazione Italiana Sindromi Neurodegenerative da Accumulo di Ferro (AISNAF, Italy), and Hoffnungsbaum e.V. (HoBa, Germany) (to L.F.B.), by the European Research Council (ERC) under the European Union’s Horizon 2020 research and innovation programme (grant agreement No. [948027]) (L.F.B.), by the Deutsche Forschungsgemeinschaft (DFG, German Research Foundation) under the Heisenberg Programme (Project No. 447395247) (to L.F.B.) and under Germany’s Excellence Strategy within the framework of the Munich Cluster of Systems Neurology (EXC 2145 SyNergy—ID 390857198) (to L.F.B. and S.F.L.) and by the Bundesministerium für Forschung, Technologie und Raumfahrt (BMFTR, FKZ: 01ED2402A) under the aegis of JPND.

We thank Philip Seibler for providing iPSC lines from two healthy control individuals (SFC156-03-012, SFC065-03-05) and from one BPAN patient (L-8172) used in this study, and the biobank “Cell line and DNA Bank of Genetic Movement Disorders and Mitochondrial Diseases,” member of the Telethon Network of Genetic Biobanks, funded by Telethon Italy, and of the EuroBioBank Network, for providing human fibroblasts from one BPAN patient (119129), and the Northwestern Stem Cell Core Facility for the subsequent generation of iPSCs. We are grateful to Deborah Kronenberg-Versteeg for her constructive feedback and helpful comments on the manuscript.

---

**Supplementary Figure 1.**
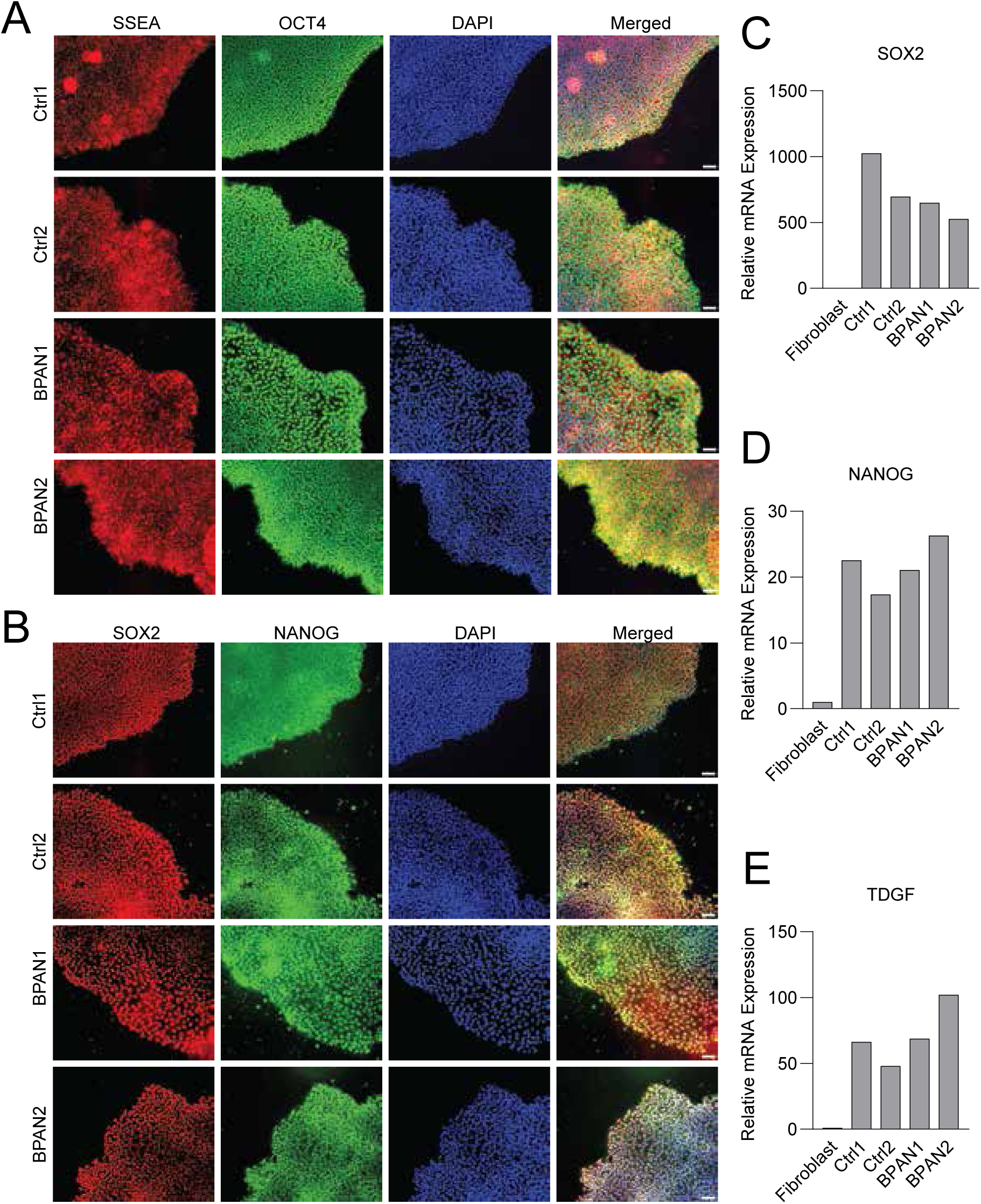
Characterization of iPSC lines from control individuals and BPAN patients. (A) Representative immunofluorescence images of pluripotency markers SSEA4 (red), OCT4 (green), and nuclear counterstain DAPI (blue) in iPSC colonies derived from two control (Control1, Control2) and two BPAN patients (BPAN1, BPAN2). Merged images are shown for each line. Scale bar, 50 μm. (B) Representative immunofluorescence images of SOX2 (red), NANOG (green), and DAPI (blue) staining in the same iPSC lines as in (A), with merged channels. Scale bar, 50 μm. (**C–E**) Quantitative PCR analysis of pluripotency marker expression in iPSC lines relative to parental fibroblasts. (C) Relative mRNA expression of *SOX2* in duplicate samples, normalized to fibroblast levels. (D) Relative *NANOG* expression. (E) Relative *TDGF* expression.

**Supplementary Figure 2.**
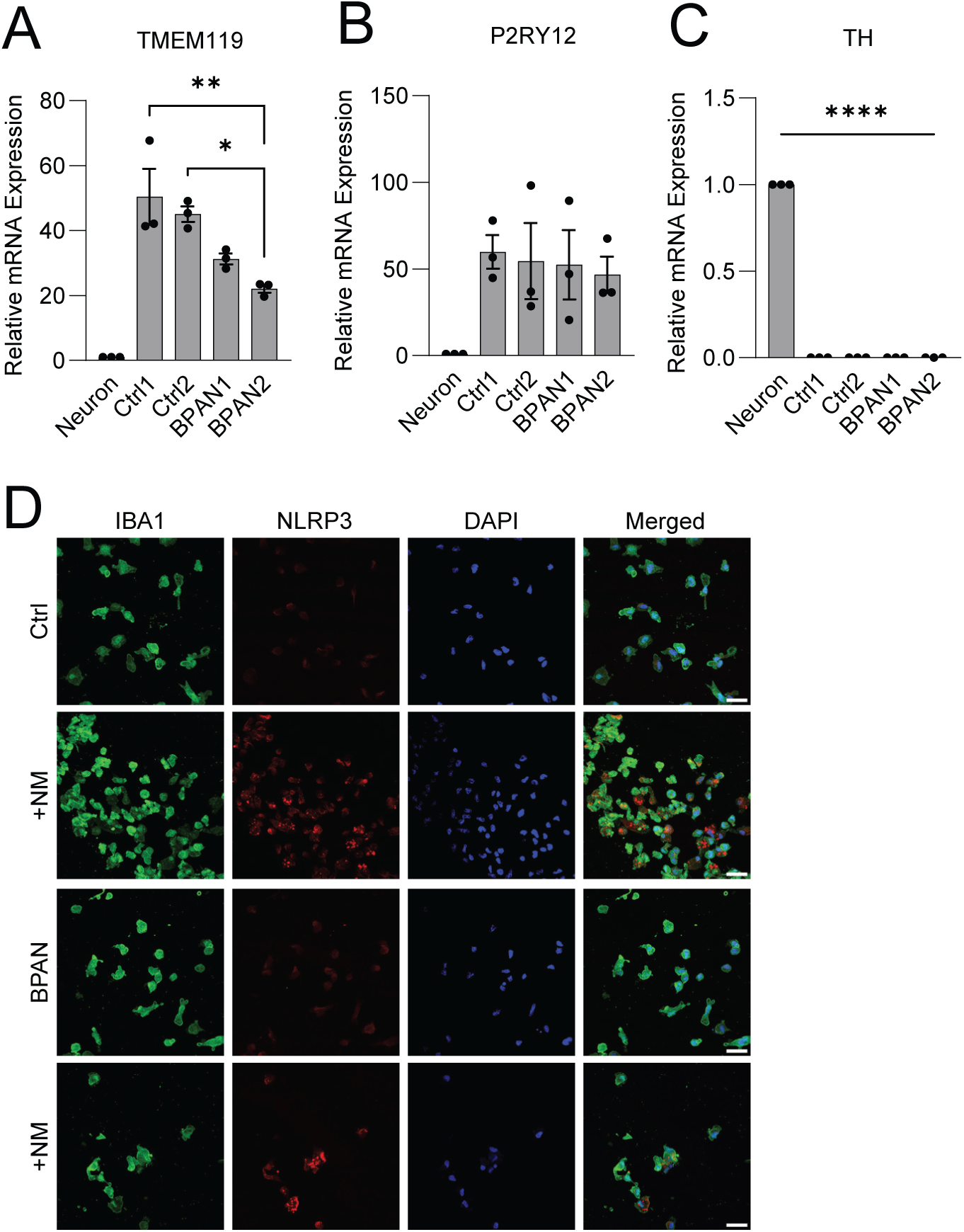
Characterization of iPSC-derived microglia culture. (**A-C**) Quantitative PCR analysis of marker gene expression in iMG compared to midbrain dopaminergic neurons. (A) *TMEM119*, (B) *P2RY12,* and (C) *TH* (tyrosine hydroxylase) mRNA levels are shown as relative expression (n = 3). (D) Representative immunofluorescence images of control and BPAN patient iMG stained for IBA1 (green), NLRP3 (red) and DAPI (blue), with and without neuromelanin treatment (5 ng/mL, 24 hours). Scale bar, 10 μm. NM, neuromelanin.

**Supplementary Figure 3.**
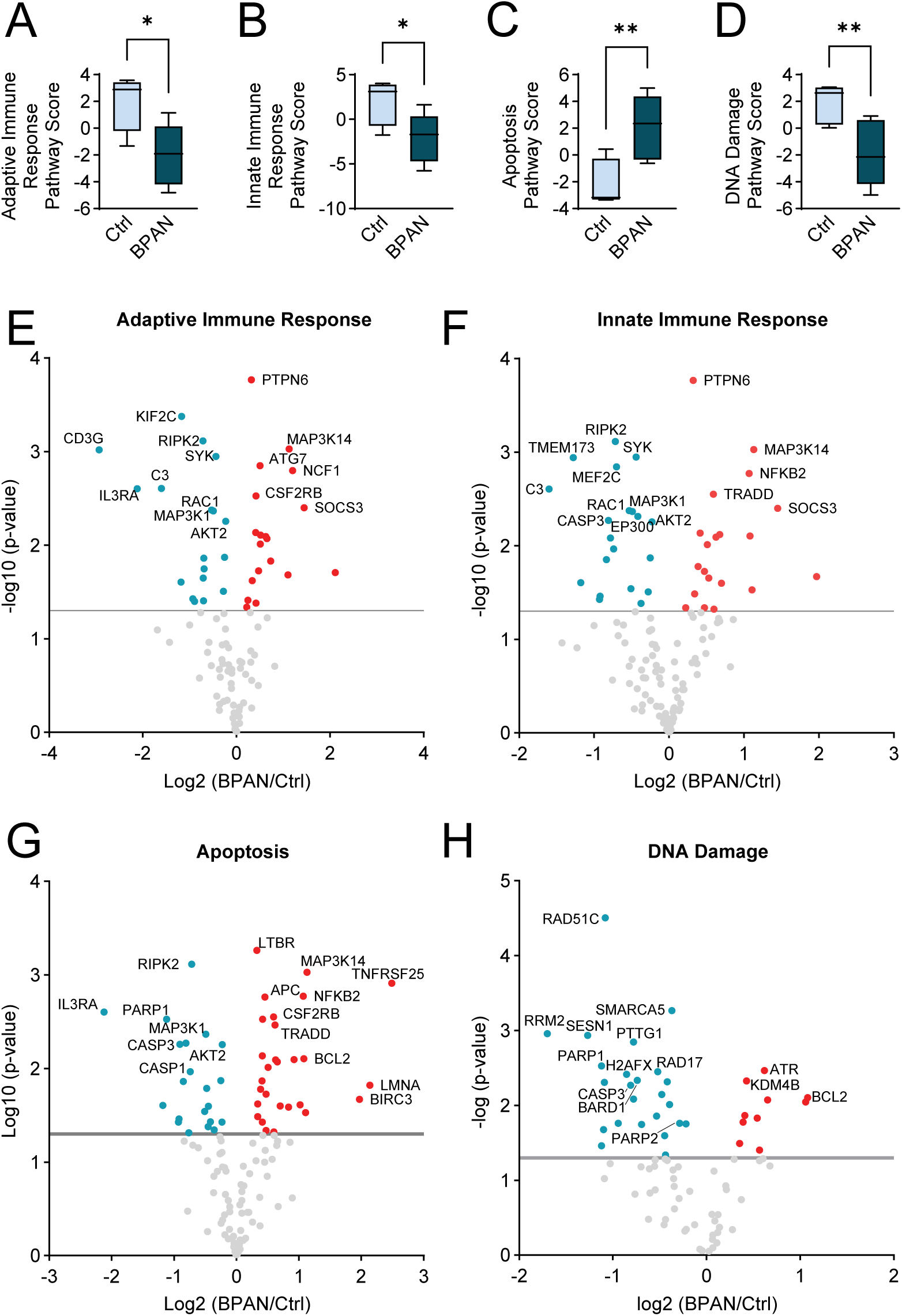
NanoString-based neuroinflammatory pathway analysis of iMG from control and BPAN patient lines. (**A-D**) Box plots showing pathway scores for selected functional categories, including innate immune response (A), adaptive immune response (B), lipid metabolism (C), DNA damage (D). Each box plot displays the distribution of pathway scores (min to max) for all samples in each group (n = 6 per group, combining 3 biological replicates from two control and two BPAN lines). Statistical significance was determined by unpaired t-test; bars represent median and interquartile range. (**E-H**) Volcano plots illustrating differential gene expression within the most relevant pathways: innate immune response (E), adaptive immune response (F), lipid metabolism (G), DNA damage (H). The top 10 genes by – log₁₀(p-value) are labeled in each plot. Red points indicate upregulated genes, blue points indicate downregulated genes, and light gray points represent nonsignificant changes. The dashed gray horizontal line marks the significance threshold (*p = 0.05*, –log₁₀ = 1.3).

